# Learning Continuous 2D Diffusion Maps from Particle Trajectories without Data Binning

**DOI:** 10.1101/2024.02.27.582378

**Authors:** Vishesh Kumar, J. Shepard Bryan, Alex Rojewski, Carlo Manzo, Steve Pressé

## Abstract

Diffusion coefficients often vary across regions, such as cellular membranes, and quantifying their variation can provide valuable insight into local membrane properties such as composition and stiffness. Toward quantifying diffusion coefficient spatial maps and uncertainties from particle tracks, we use a Bayesian method and place Gaussian Process (GP) Priors on the maps. For the sake of computational efficiency, we leverage inducing point methods on GPs arising from the mathematical structure of the data giving rise to non-conjugate likelihood-prior pairs. We analyze both synthetic data, where ground truth is known, as well as data drawn from live-cell singlemolecule imaging of membrane proteins. The resulting tool provides an unsupervised method to rigorously map diffusion coefficients continuously across membranes without data binning.

## 1 Introduction

Cellular membranes play critical roles in many important biological processes, such as signal transduction [1], molecular transport, and the maintenance of structural integrity of cells [2]. Owing to the complexity of cellular membrane architecture and its interactions with peripheral structures on both the interior and exterior of the cell [3], modeling membrane and, more generally, heterogeneous protein diffusive dynamics in space remains an active area of study [4, 5, 6, 7].

To understand the dynamics of embedded membrane proteins, several methods have been proposed to map diffusion coefficients, or diffusivities, of membrane proteins [8, 9, 10, 11, 12, 13, 14].

While all of these methods infer dynamical quantities, most of them involve data binning as a data pre-processing step. This necessarily reduces the amount of information left to analyze in deducing diffusion coefficient maps whether from membrane proteins or other applications. Beyond data reduction, binning methods, ultimately, impact the spatial precision of inference and our ability to rigorously propagate error arising from particle localizations into diffusion coefficient spatial maps. Approximately optimizing bin sizes, locations, and shapes in estimating diffusion coefficient magnitudes has been addressed, whether involving Voronoi tessellation [15] or assigning length scales to each localization based on noise [16], though leaving unresolved the possibility of avoiding data binning altogether.

Furthermore, inherent to binned analysis is the implication that diffusion coefficients are assumed to vary across bins, but otherwise remain constant within any one bin [13]. This sharp spatial variation, introduced by binning, masks the precise underlying gradient of the diffusion coefficient change within a bin that may already be encoded in the data. Recent approaches, based on deep learning [14], remove the need for binning but require supervised training on labeled datasets.

To address these fundamental issues, we propose a new framework circumventing these difficulties by determining a continuous diffusion coefficient map using all available data without binning in an unsupervised fashion. That is, we are focused on learning continuous 2D diffusion coefficient maps from trajectory data. In principle, the trajectory data can be of any type and the spatially varying diffusion coefficient can arise from any number of physical origins.

To achieve this, we develop a Bayesian framework that uses trajectory data to infer all candidate diffusion coefficient maps alongside their associated uncertainty. Within our Bayesian framework, we avoid data binning by leveraging the mathematics of Gaussian Process (GP) priors on the continuously defined diffusion map that we wish to learn. We further leverage approximations (inducing point methods detailed later [17, 18]) to otherwise reduce the cubic scaling [19] of naive GP implementations.

We demonstrate our method on both synthetic data, where ground truth diffusion maps are known, as well as experimental data involving membrane protein trajectories extracted from live-cell single-molecule imaging experiments on cellular membranes.

## 2 Methods

Our goal is to infer continuous spatial diffusion coefficient maps in 2D, from particle trajectory data. In many practical examples, these particles are membrane proteins. Then we provide a numerical scheme suited for the method. Finally, we validate our method on both synthetic and experimental data.

### 2.1 Theory Methods

Concretely, we treat diffusion dynamics as a Brownian random walk, under the Itôo approximation [20, 21], with spatially varying diffusion coefficient. Under this approximation, we first describe a forward model relating the collected data to spatial diffusion coefficient maps. Next, we design an inverse method, to learn diffusion coefficient landscapes warranted by the data. Strictly speaking, as we work within a Bayesian paradigm, we develop a posterior over all candidate spatial maps to which we can assign probabilities. We then use Monte Carlo to sample from our posterior. Though in principle our framework assigns probabilities to all putative spatial maps given the data, for convenience, the spatial maps we illustrate in all figures are those maximizing the posterior termed the maximum *a posteriori* (MAP) 2D maps.

#### 2.1.1 Forward model

We index each particle, to which we associated one track (*i*.*e*., one location in each frame) using *i* = 1, .., *I*, with *I* the total number of particles. For each particle, location measurements occur at fixed time intervals, spaced Δ*t* apart, defining time frames. Not all particles are present in each frame. To accommodate this contingency, we define a quantity *N*_*i*_ which is the total number of frames in which particle *i* appears. For instance, particle 1 (*i* = 1), may appear in a total of 2 frames, and thus *N*_1_ = 2. Particles are assumed to appear in consecutive frames and, if they disappear due to blinking of the labels on membrane proteins for instance in fluorescence experiments, then we consider the particle once re-appeared as a new particle. As we are only learning diffusion coefficient maps, and not keeping track of particle identities, this convention is purely a matter of convenience.

Next, we define the position of a given particle, *i*, in its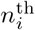 frame as 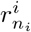. Within a free diffusion model with spatially varying diffusion coefficient, we assume this position to be normally distributed around the preceding position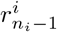, with variance proportional to the diffusion coefficient at that previous position. This transition probability is expressed as

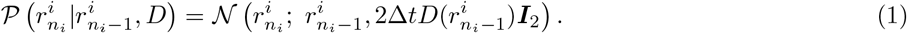

Here *D*(*·*) is a continuous function describing the spatially dependent diffusion coefficient and ***I***_2_ is a 2D identity matrix. Using the transition probability previously defined, we construct the likelihood of a collection of *I* trajectories with *N*_*i*_ positions for each *i* particle, given a diffusion coefficient map, *D*, as follows

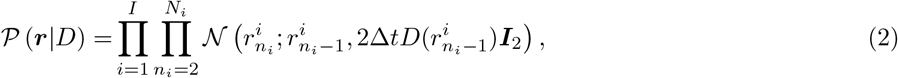

where 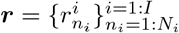 collects all observed particle positions for all particles.

#### 2.1.2 Inverse method

Given our forward model, our next task is to develop the (posterior) probability distribution over diffusion coefficient maps. For this, we use Bayesian inference, as it provides a principled framework to systematically incorporate observed data, leading to reliable and robust estimation. In mathematical terms, this is achieved leveraging Bayes’ Theorem

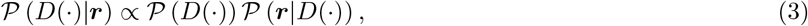

where 𝒫 (*D* (*·*)) is the prior distribution on all candidate maps, 𝒫 (***r***| *D*(*·*)) is the likelihood of the data provided a specific diffusion coefficient map, given by Equation (2), and 𝒫 (*D*(*·*)| ***r***) is the posterior distribution assigning probabilities to all candidate diffusion coefficient maps given the data.

Having specified the likelihood in Equation (2), we now specify a prior. Specifically, we need a distribution on *D*(*·*) assigning probabilities to continuous surfaces but also allowing for a convenient form when evaluated at discrete spatial positions. A common choice is the Gaussian Process (GP) prior [19, 18]. GPs are an infinite collection of co-varying random variables, any finite subsample of which follows a multivariate Gaussian distribution, allowing for a means by which to assign a probability over function space. In this case, the function space consists of surfaces. This means that we can assign probabilities to continuous diffusion coefficient maps based on a discrete set of values.

Thus far, we have been using *D*(*·*) to refer to the continuous diffusion coefficient function. As we begin to explicitly define the inference task, we will need to define this function on a 2D discrete grid as finely spaced as computational efficiency permits. Notationally, we use ***D*** to represent an array of diffusion coefficients and use an appropriate subscripted index to describe the locations of the discretization. Selecting a finite number of training points, *ω*, on the diffusion coefficient surface, the prior approximates to

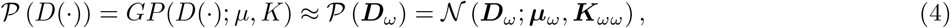

here ***D***_*ω*_ is the array of diffusion coefficients based on an arbitrary surface at locations *ω*, ***μ***_*ω*_ is the mean of the prior at those same training points, and ***K***_*ωω*_ is the auto-covariance between the training points.

Evaluating the posterior typically involves inverting the covariance matrix [19] which becomes computationally expensive, scaling cubically with the number of data points [18], and unstable for large datasets. To address this, we turn to an inducing point method [17] defining a uniform grid of points, performing inference on those points, and obtaining a finer resolution when needed by leveraging a covariance-based interpolation method. For our inducing point model, we establish *m* inducing points as a uniform grid on the domain on which we infer ***D***_*m*_. That is

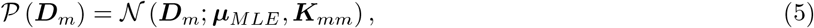

where ***D***_*m*_ is the array of diffusion coefficients at the inducing points, ***μ***_*MLE*_ is the mean of the prior, and ***K***_*mm*_ is the auto-covariance matrix between the inducing points. Having defined a computationally efficient form of our prior, we can proceed by defining values for the two hyperparameter quantities, mean and covariance. As a convenient starting point, we set the mean of the prior to be a flat diffusion coefficient with magnitude given by the Maximum Likelihood Estimate (MLE) over the entire data [22],

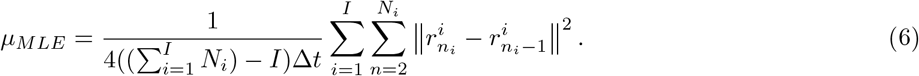

As for the covariance we choose a square exponential kernel [18], also known as the Gaussian Radial Basis Function (RBF) whose form reads

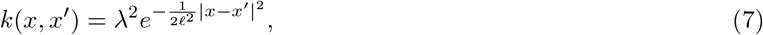

here *λ* sets the variance of the multivariate Gaussian and *ℒ* defines the covariance between positions *x* and *x*^*′*^. The explicit values of these hyperparameters can be tuned based on the expected variation of the true diffusion surface. Generally, to keep the prior uninformative, we set the variance, *λ*, to be twice the magnitude of the MLE, and set the length scale to 20% of the data’s range.

For computational convenience, we placed the prior on a coarser, inducing point, grid though our likelihood is on the grid of available (unprocessed, *i*.*e*., unbinned) data points. Thus to compute both prior and likelihood simultaneously, we must have a way to interpolate from one grid to the other. This is achieved by rigorously interpolating using our covariance matrices [17, 18]

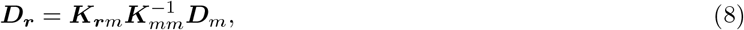

whose elements read

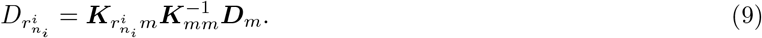

Here 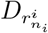 is the diffusion coefficient at 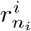 and 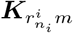 is a row vector, from the full covariance matrix ***K***_***r****m*_, whose elements are the covariance between the position 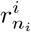 and all inducing points.

Under our inducing point interpolation scheme, we reparameterize our likelihood, Equation (2),

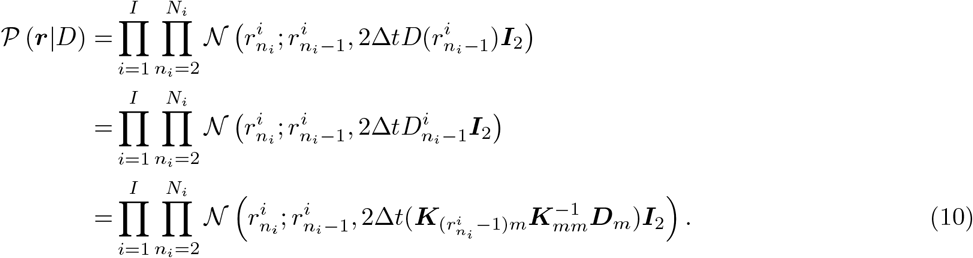

This setup yields the following posterior distribution (up to a normalization constant) over ***D***_*m*_ given the data, ***r***,

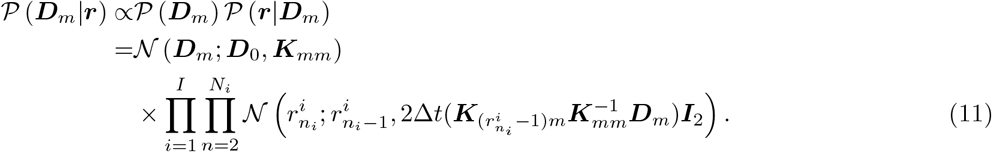

By maximizing this posterior with respect to ***D***_*m*_, we obtain the most probable diffusion coefficient distribution explaining the observed data ***r*** though sampling from the posterior is possible to gain information on uncertainty. While direct sampling from this posterior is challenging due to the mathematical (non-conjugate) form of the prior and likelihood, we rely on MCMC sampling [18], specifically by constructing a Metropolis-within-Gibbs scheme.

#### 2.1.3 Algorithm

Metropolis-within-Gibbs sampling requires a proposal distribution for generating samples of ***D***_*m*_. A straightforward approach is to propose new 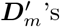, either element-wise or as whole surfaces, based on the previous value of ***D***_*m*_ [23]. However, such a naive approach is problematic because the prior, from Equation (5), favors small deviations from the mean. This necessitates proposals amounting to small variations away from ***D***_*m*_, leading to extended convergence times with no guarantee of avoiding local optima traps.

To address this challenge, we introduce a reparametrization that allows us to propose new values for ***D***_*m*_ in a more efficient manner. We linearly transform the array of inducing points, using the inverse auto-covariance matrix, into a space that respects the covariance of the prior, denoted as 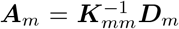. In this transformed space, we can make proposals across the entire map whose smoothness matches, and is therefore less penalized, by the prior enabling more substantial proposals.

To expedite convergence, we initialize the Metropolis-Hastings algorithm at a highly probable sample deterministically identified. In our specific case, a very effective initialization method involves performing MLE at the data points with interpolation to the inducing points using RBF interpolation. For this, we use the same RBF as our covariance matrix Equation (7).

The last challenge with a high dimensional posterior is avoiding local posterior maxima, while the algorithm is iteratively converging. We tackle this in two ways, the first is tempered sampling [24], which allows us to transform our posterior into an augmented space where we can control the behavior of the sampler. Mathematically we write

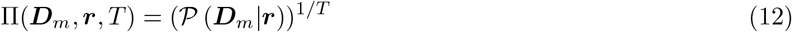

where the temperature (*T*) of the sampler dictates how heavily the sampler is penalized for moving to areas of low probability. Sampling at higher temperatures results in a lower penalty and vice-versa, while a temperature of 1 is precisely our standard Monte Carlo scheme. For our specific case, we begin with a temperature of 10 and exponentially decay to 1.

The second way we avoid local maxima is by stochastically iterating through the components of ***A***_*m*_ when making proposals. This means that instead of proposing new values of the components in order every time, we iterate through the components of ***A***_*m*_ randomly. This prevents proposals in any one area of the surface from dominating the accepted samples.

With these modifications, we outline the algorithm as follows.

##### Initialization

- We first define our hyper-parameters, *μ*_*MLE*_, *λ, ℒ*, as outlined in Equation (6) and Equation (7).
- Compute the initial sample deterministically using the MLE at each point and performing a kernel-based interpolation, using Equation (7) as the kernel, to the inducing points 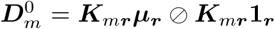. Here ⊘ is Hadamard division (elementwise division).
- Transform the initial sample into inverse space: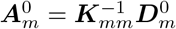

##### Monte Carlo

- For many iterations, perform the following steps: Begin with the desired sampling temperature, 10 in our case, and allow it to decay to 1 over the iterations. During the decay, generate samples of ***D***_*m*_ using 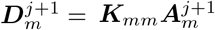 is obtained through stochastic iterations over *k*, the inducing points in inverse space:

1. Propose new values for 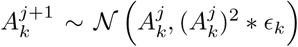, where 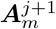 is a constant for adjusting the proposal magnitude to maintain a desirable acceptance rate (about 25%). *ϵ*_*k*_ is automatically updated after every full sweep through *k*.
2. Accept or reject the proposal based on the tempered Metropolis-Hastings acceptance ratio while adjusting for the current temperature of the sampler. One full loop through all values of *k* is considered the new sample 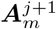.

##### Post-Processing

- After the sampler has run and converged, select the sample with the highest probability to obtain the MAP estimate.

This algorithm efficiently explores the high-dimensional posterior over the diffusion map and also estimates the most probable diffusion coefficient map (the MAP estimate) given the observed data.

### 2.2 Data Methods

#### 2.2.1 Generating and Benchmarking Synthetic Data

To benchmark our inference method and assess its performance across various scenarios, we initially rely on the use of synthetic data. Synthetic data provides a controlled environment for rigorous evaluation, allowing us to precisely control the properties of the data and establish known ground truth diffusion coefficients. By comparing the algorithm’s estimated diffusion coefficient maps with these known ground truths, we validate its effectiveness and identify potential limitations. Synthetic data are generated by specifying an arbitrary diffusion coefficient map surface, and then simulating trajectories by directly sampling from the forward model, specifically Equation (2).

In our evaluation, we introduce specific challenges to test the algorithm’s capabilities:

1) Reducing the amount of data. To evaluate how well the algorithm performs under conditions of limited data availability, we systematically reduce the amount of data in our synthetic scenarios. This allows us to benchmark the method’s accuracy with decreasing data. More concretely, we simulate more challenging data scenarios by reducing the number of particles or randomly dropping positions from particle trajectories. This helps us replicate situations where certain regions have few measurements or the particle is lost over tracking, say, due to phenomena such as photobleaching. Representative results are shown in Figures 2a, 2b, 2c, while additional full analyses are available in Figure S1;

**Figure 1:**
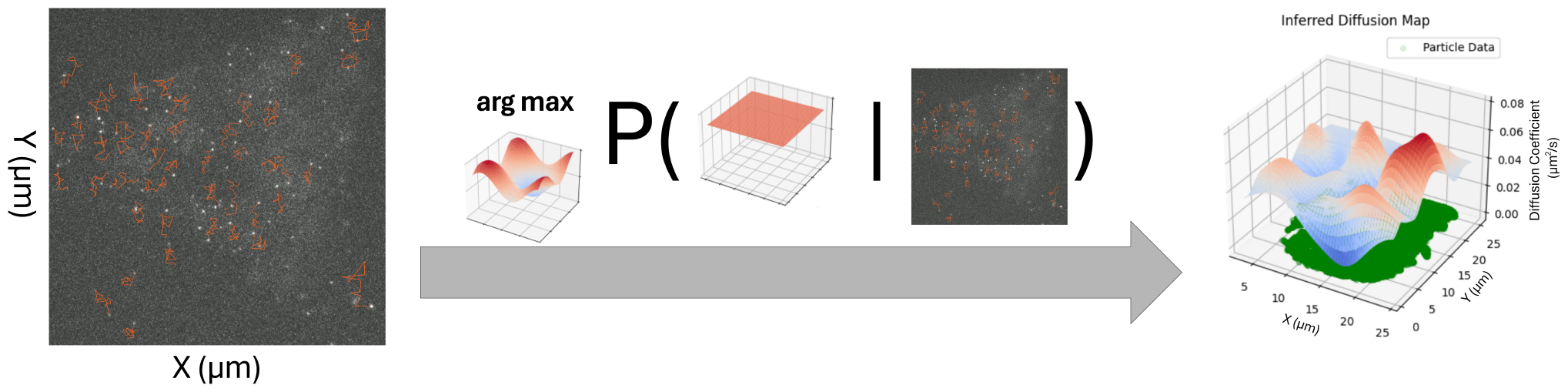
Schematic representation of our method. Our method uses single-particle localizations forming trajectories as input and outputs a continuous surface describing the diffusion coefficient as a function of space without binning or other forms of data downsampling. On the left is a single frame from a fluorescence microscopy frame stack which has been artistically labelled in orange to represent membrane protein trajectories. These trajectories are the input into our model, which identifies the spatial diffusion coefficient map of the highest probability. This is plotted on the right, with green dots identifying the localizations of particles used in deducing the diffusion coefficient map. The linking between the localizations of each particle across each frame forms tracks which, for clarity alone, are not shown.

**Figure 2:**
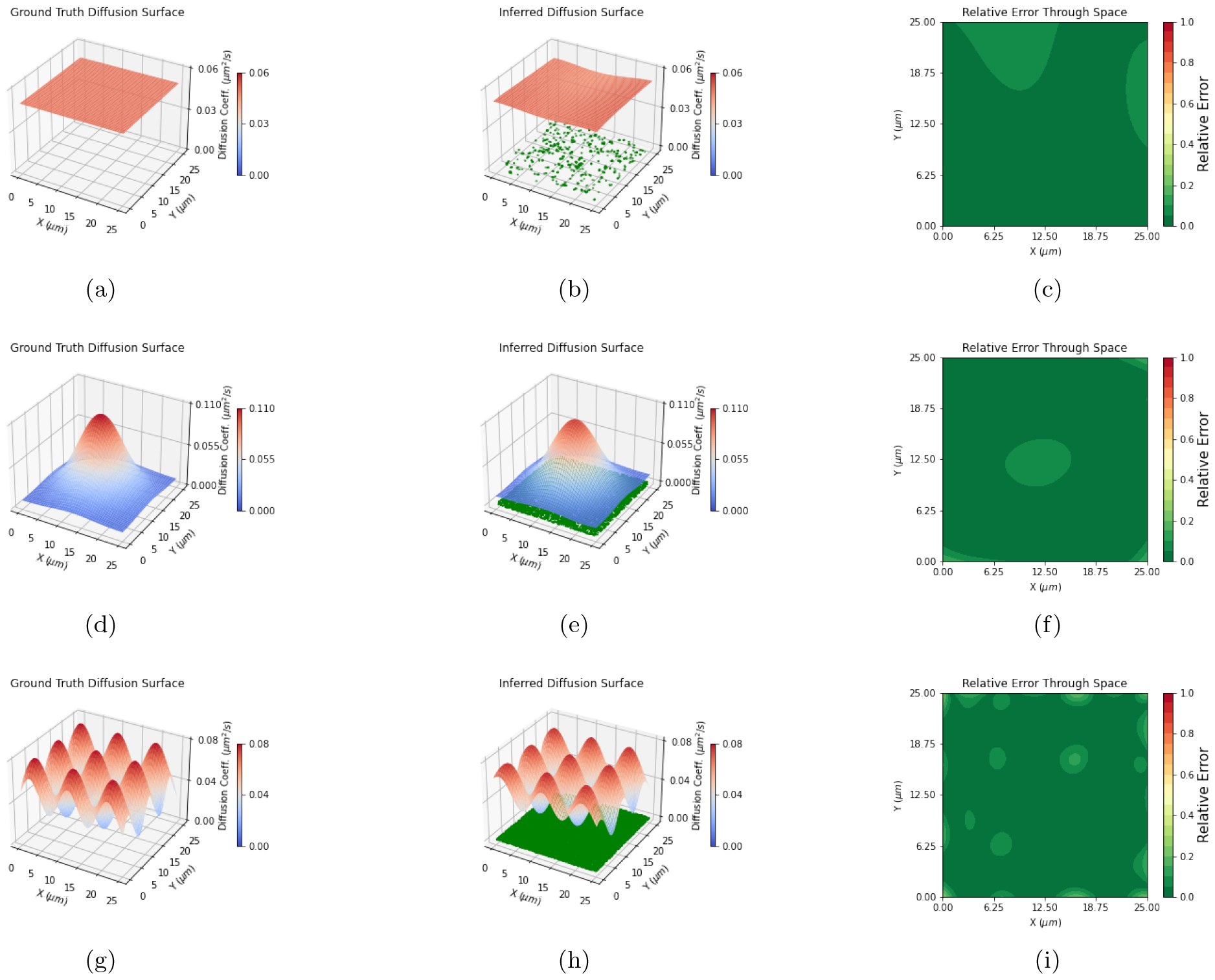
Learning diffusion coefficient maps from synthetic data. Each row here represents an analysis of a unique synthetic data set. The first column of each row shows the true diffusion coefficient map used in synthetic data simulation and the second column plots the diffusion coefficient map inferred by the algorithm, with the synthetic data plotted in green below the surface. As can be seen in Figures 2b 2e 2h, we progressively increase the number of localizations, 5 *×* 10^3^, 10^5^, and 2 *×* 10^5^ respectively. The third column plots the relative error between the Ground Truth Diffusion Map and the Inferred Diffusion Map as a function of space computed according to Equation (13)

2) Increasing the complexity of the diffusion coefficient maps used in generating the synthetic data. That is, to assess the algorithm’s performance in the presence of important spatial changes in diffusion coefficients, we create synthetic data scenarios with rapidly spatially varying diffusion coefficients. For instance, we generate diffusion coefficient maps as wave patterns with progressively decreasing wave periods in each dataset to test the method’s robustness. By generating synthetic data under these conditions, we gain valuable insights into the algorithm’s behavior, ensuring its robustness and adaptability across a range of practical situations. Representative results are shown in Figures 2g, 2h, 2i, while additional full analyses are available in Figure S1.

Once we have generated and analyzed such synthetic datasets with our algorithm, we benchmark the results with a calculation of spatial accuracy using:

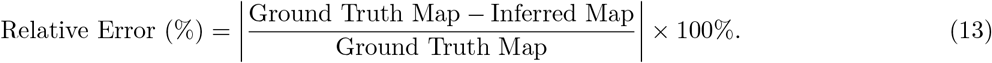

#### 2.2.2 Experimental Methods

For the live-cell single-molecule imaging experiments, CHO cell lines stably expressing dendritic cell-specific intercellular adhesion molecule-3-grabbing nonintegrin (DC-SIGN) wild-type and DC-SIGN N80A mutant, established by Lipofectamin 2000 (Invitrogen) transfection, were cultured in Ham’s F-12 medium (LabClinics), supplemented with 10% heat-in-activated FBS (Invitrogen), 1% Antibiotic Antimycotic Solution (GE Healthcare Life Sciences), and 0.5 mg/mL of the aminoglycoside antibiotic G418 (Invitrogen). For the protein labeling, we used half-antibody fragments obtained following a protocol similar to the one used in [25]. DCN46 antibody (Pharmingen) was dialyzed overnight at 4ôoC using Slide-A-Lyzer MINI Dialysis Units (Thermo Scientific) in PBS and reduced with DTT (Invitrogen) following the manufacturer’s instructions. Reduced antibodies were then biotinylated with Maleimide-PEG2Biotin (Thermo Scientific). Nonreacted DTT and unbound biotin were removed by overnight dialysis. Biotinylated half-antibody fragments were conjugated with streptavidin-coated quantum dots QD655 and QD585 (Invitrogen). Before imaging, CHO cells were seeded onto 25-mm glass coverslips (Menzel-Gla asser). Cells were incubated with quantum dot conjugates for 5 min at RT. Extensive washing with serum-free medium was performed to remove unbound conjugates. Imaging was performed using an Olympus fluorescence microscope equipped with a 1.4 NA, 100 *×* objective. Samples were illuminated in epifluorescence geometry with the 488-nm line of an argon-ion laser (Spectra Physics), with power density at the sample plane of approximately 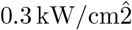. Emission light was split with appropriate dichroic mirror and filters, and collected on an intensified EM-CCD (Hamamatsu). Movies were recorded at a frame rate of 33 Hz for 10,000 frames. Detection and tracking were performed using u-track [26]. The detection (Gaussian Mixture-Model Fitting) and tracking parameters were optimized based on visual inspection and performance diagnostic of the resulting detection and tracking. All image and data analysis were performed in MATLAB (The MathWorks, Natick, MA). Videos were loaded into MATLAB using Bio-Formats [27].

## 3 Results

As mentioned previously, we split the discussion of our results into two sections. The first pertains to results on synthetic data used to validate the robustness of our algorithm. The second section consists of experimental data to highlight our framework’s applicability to biological data. Our standard for an accurate map will be a 10% relative error, *i*.*e*., as long as we stay below 10% we claim that we have inferred the diffusion map accurately.

### 3.1 Synthetic Data

Here we show the results on three unique synthetic datasets. For all synthetic datasets, we keep a consistent 25-micron square for the field of view and randomize both the exact number of trajectories and the length of each. Each dataset validates our inference algorithm and showcases an important feature of the method. The first, Figures 2a, 2b, 2c, shows our inference task on a flat diffusion surface, showing how inferring a continuous surface is possible within our chosen Bayesian paradigm using only about 5 *×* 10^3^ localizations (from around 250 trajectories). In Figures 2d, 2e, 2f, we show that it is possible to learn a single perturbation upon the flat surface, verifying that the model is able to infer accurate diffusion maps using about 10^5^ localizations. Finally, simulate the data using around 2 *×* 10^5^ localizations using a diffusion coefficient map consisting of a series of waves where the maximum period is 40% of the field of view. In Figures 2g 2h 2i, we see that we can recover the diffusion map within 10% relative error with few areas just outside that threshold.

### 3.2 Experimental Data

Here we show results obtained for experimental data of live-cell single-molecule imaging. Dendritic cell-specific intercellular adhesion molecule-3-grabbing nonintegrin (DC-SIGN) stably expressed in CHO cells were fluorescently labeled and imaged on the dorsal membrane of living cells. Previous work [28, 29] has reported heterogeneous diffusion for both the wild-type and N80A mutant, characterized by changes in diffusion coefficient.

For all experimental data, the field of view is under or equal to a 25-micron square, and the datasets themselves contain at least 1.8 *×* 10^5^ protein localizations. After running on synthetic data for parameter regimes close to this (and showing the method infers diffusion coefficient maps within 10% relative error, see Figure 2), we now turn to experimental data.

The results in Figure 3 show that we are able to recover diffusion coefficient maps from the various sets of membrane protein trajectories. The magnitude of the diffusion coefficient recovered by our method is in line with previous observations [28, 29], shown in Figure S3. As always, with experimental data, we do not have a ground truth to assess the accuracy of our inference.

**Figure 3:**
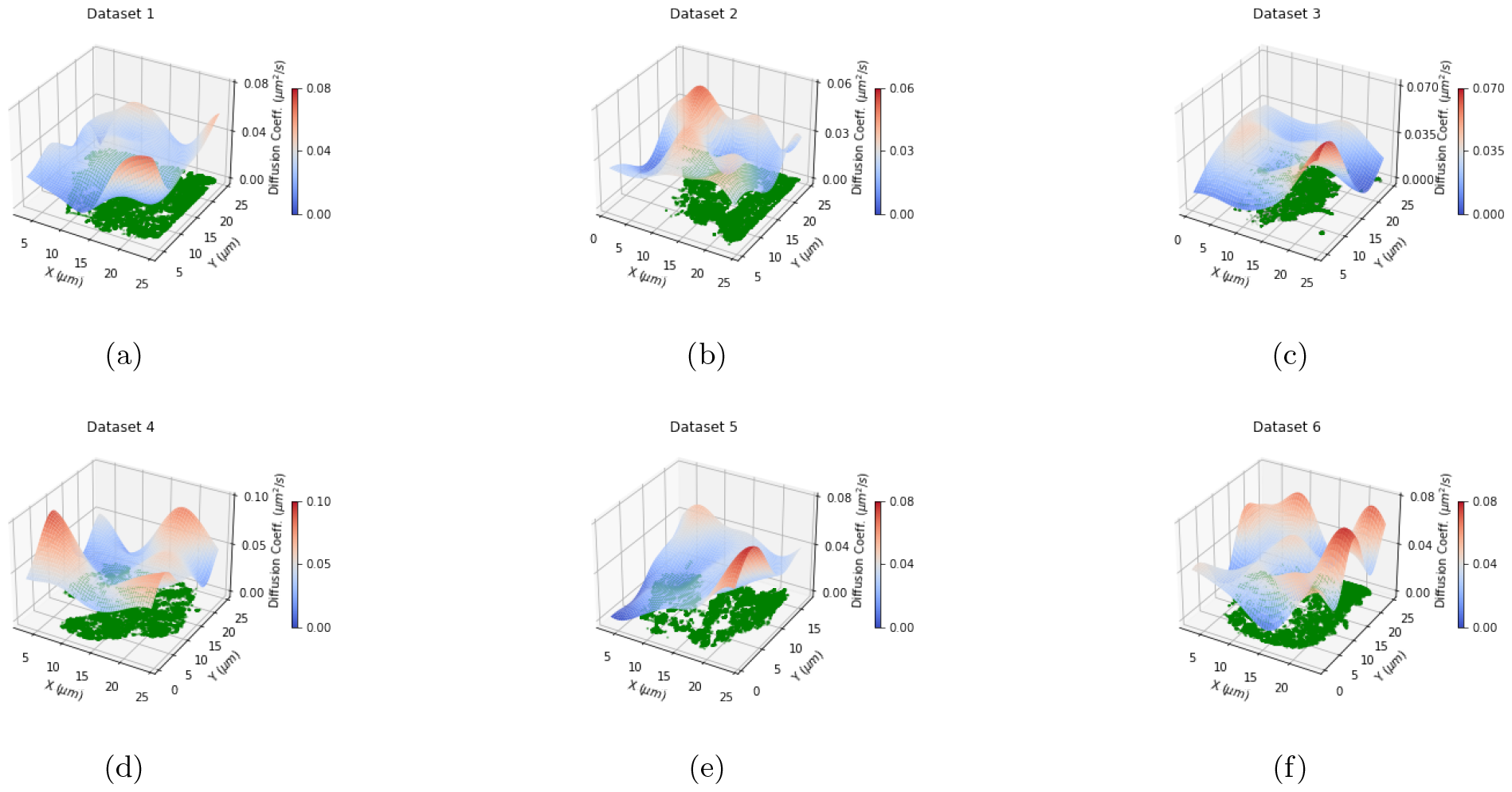
Learning diffusion coefficient maps from experimental data. Here we visualize the inferred diffusion coefficient map from six different experimental datasets, each from different cells. The green points at the bottom of each plot represent DC-SIGN wt (a-c) and N80A (d-f) 10^5^ localizations from trajectories analyzed for each set. The surfaces plotted are the inferred surfaces from the algorithm and, as expected, they diverge towards the edge where there is no data as we intentionally analyze a region larger than the data provided to extrapolate the diffusion coefficient map slightly beyond the data.

To address this, we produced a self-consistency check by splitting all experimental datasets in half and running the algorithm separately on each half to verify the convergence of both to a consistent diffusion coefficient map. To maintain similar spatial densities, we split the data into two subsets by extracting alternate positions from each protein trajectory. Figure 4 shows the results after subsetting the data. Here we have purposefully extrapolated beyond areas of data for two reasons: the first is for visualization convenience and the second is to analyze behavior slightly beyond the data. Allowing for the fact that GP inference on diffusion maps tends to revert to the prior at the edges with limited data [30], we see that from the remaining areas with data, the inferred maps derived from each subset stay within 10% relative error, the same standard used for the synthetic data. In addition, in SI we split the data by randomly selecting trajectories and comparing the diffusion maps from both data subsets demonstrating that the diffusion map is constant in time within error over the observation time, shown in Figure S2.

**Figure 4:**
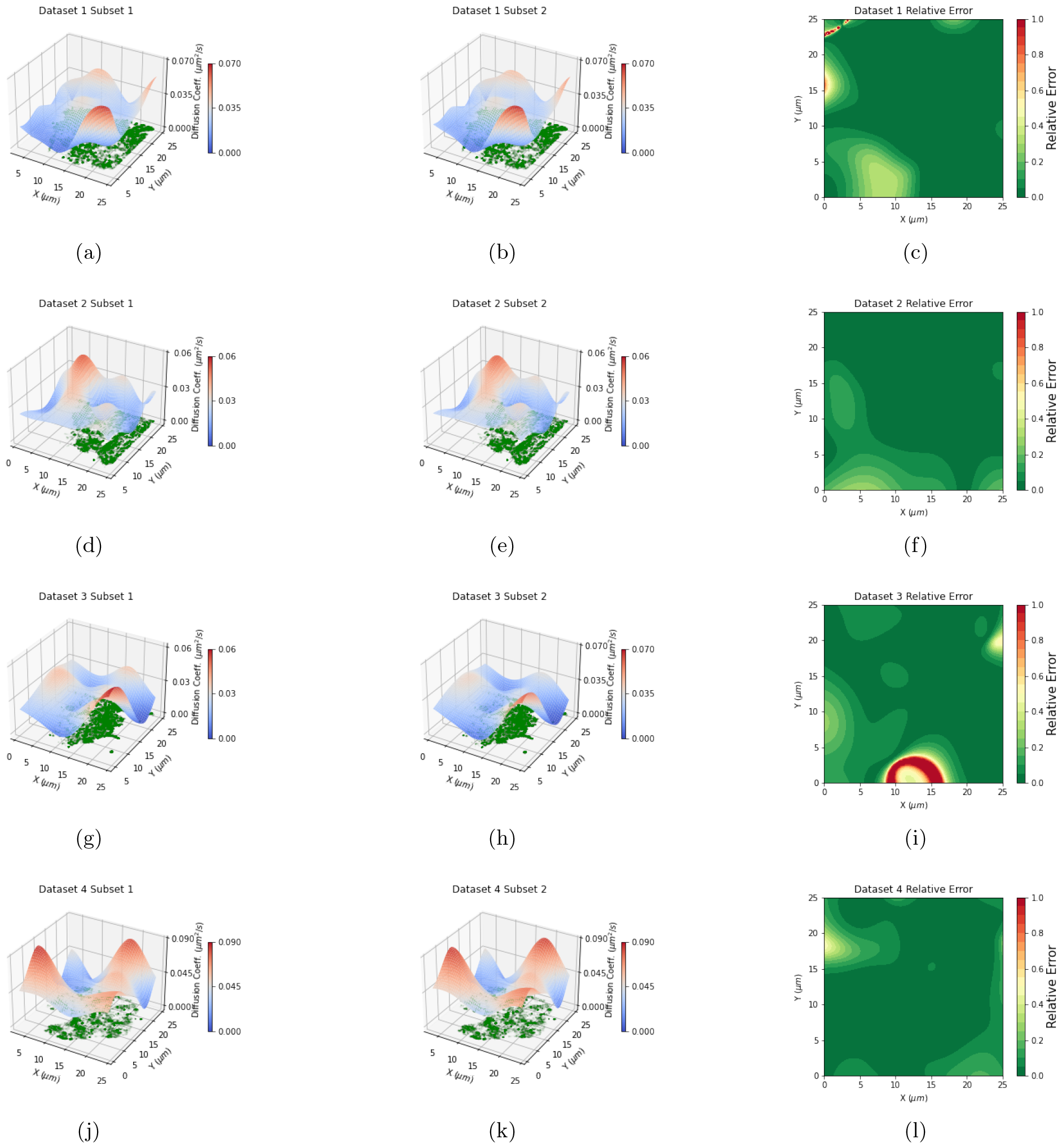
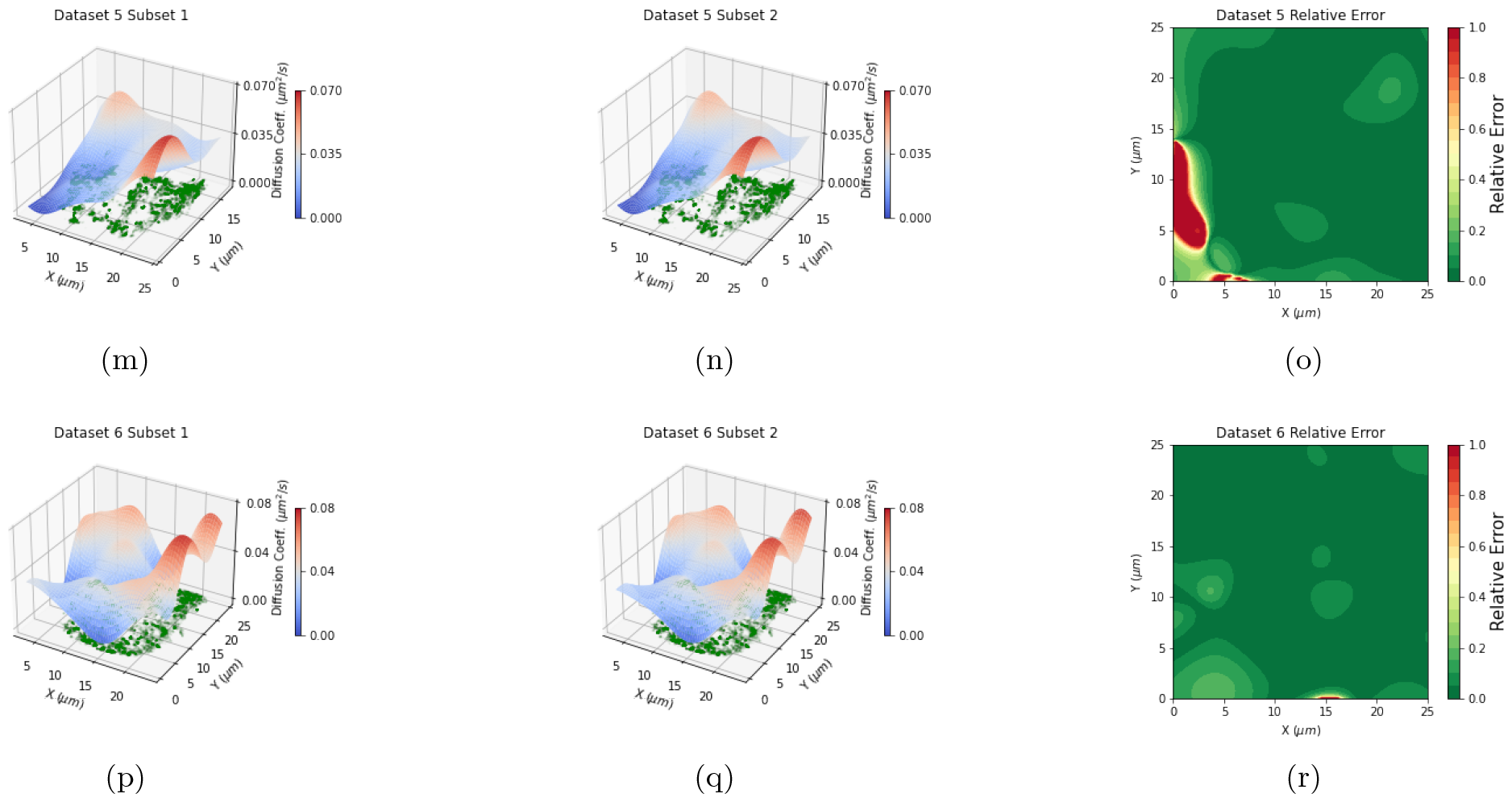
Self-consistency check on experimental data. We subsetted each experimental dataset into half and ran the algorithm on each half. Each row above represents a unique dataset. The first two columns coincide with the inferred surface plotted with the respective data half below. The third column is the relative error between the two inferred surfaces. In areas of the membrane that have protein trajectories, we can see that the relative error between the two surfaces stays below 10%. We also highlight that the red in Figures 4c 4i 4o 4r specifically arises in areas where no data is available and thus differences from samples from the very broad prior in both MAP estimates are very different.

## 4 Discussion

We have presented a general inference algorithm that can accurately infer continuous spatial diffusion coefficient maps from particle trajectories. Working within a Bayesian framework, we developed a posterior distribution assigning probabilities to all possible diffusion coefficient surfaces. We were able to reduce the computational burden of naive GP regression by adopting an inducing point method [17] auxiliary variable sampling techniques. More concretely, we reduced the otherwise cubic scaling of naive GP with the data [19, 17, 18], 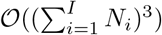 to 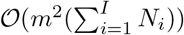 for initialization, 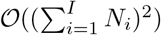 to 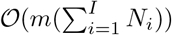 per Monte Carlo iteration, substantially reducing the computational burden and enhancing the efficiency of the posterior sampling.

We analyzed the accuracy and robustness of our method through the analysis of synthetic data, with a known ground truth. The results show that our model successfully captures the spatial variation of the diffusion coefficient, with relative errors within 10%. This validates the reliability of our approach. Furthermore, we have applied our method to experimental data verifying self-consistency within 10% relative error by splitting the data into two halves and analyzing each independently.

Ultimately, moving forward, improved uncertainty quantification in the diffusion coefficient maps recovered may be achieved by using tools capable of quantifying localization uncertainty [31, 32] and propagating this uncertainty into an uncertainty over the diffusion coefficient map. However ideally, at computational cost, we ambitiously envision future work simultaneously and self-consistently learning diffusion coefficient maps while tracking. In other words, it may be possible to envision a more general framework avoiding the modular structure proposed here that first requires tracks as input and then processes these tracks to produce diffusion maps. This modular structure is acceptable for well separated tracks but may start to fail for crowded regions [31, 32] with many particles criss-crossing paths.

Such a self-consistent framework avoiding modularity may also benefit the analysis of dimmer particles to which larger localization uncertainty is associated and treat other sources of heterogeneity in the biological data. For example, it may shed quantitative insight on the role of lipid composition on protein diffusion by correlating lipid localization to diffusion maps [33, 34, 35].

## 5 Code Availability

The code for this work can be found at https://github.com/LabPresse/GPDiffusionMapping

## 6 Acknowledgements

We thank Maria Garcia-Parajo for providing data collected in her lab. We acknowledge the support of the NIH (NIGMS R01GM130745, NIGMS R01GM134426, NIGMS R35GM148237). C.M. acknowledges support through grant PID2021-125386NB-I00 funded by MCIN/AEI/10.13039/501100011033/ and “ERDF A way of making Europe”.

## Supplemental Information

### S0.1 Robustness on Synthetic Data

**Figure S1:**
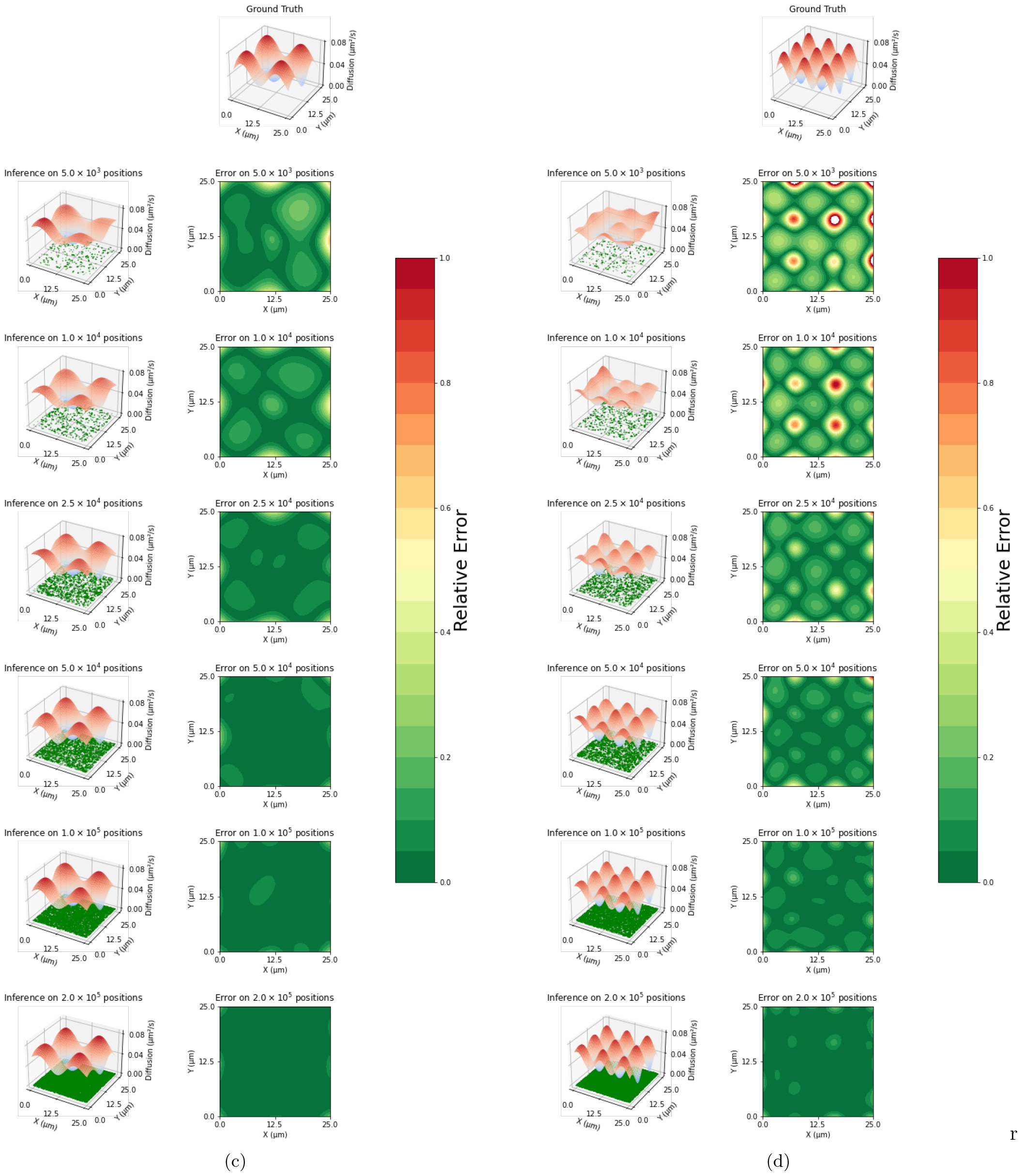
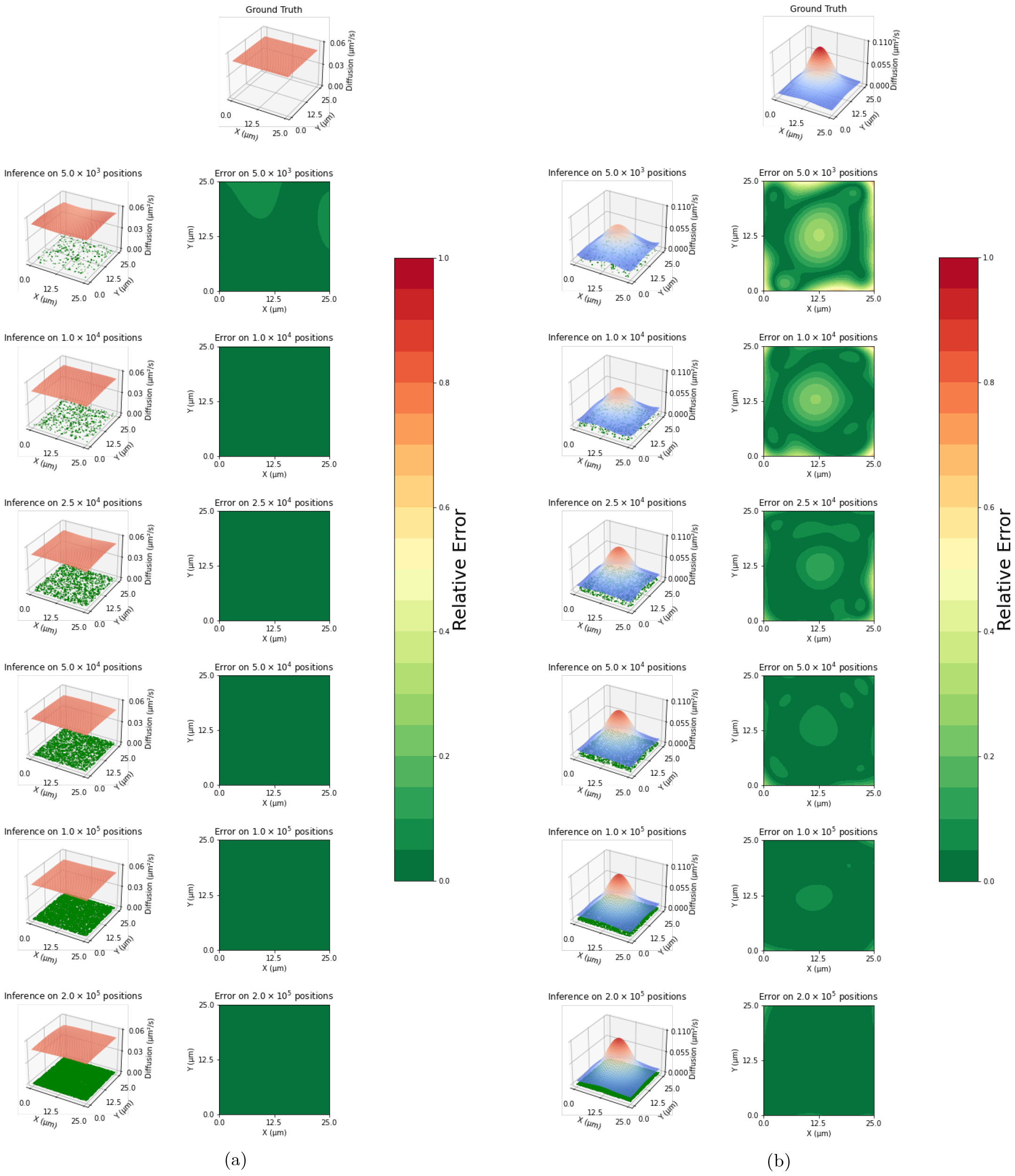
Robustness of inference on synthetic data. Here we evaluate our model’s robustness with respect to varying diffusion maps and the amount of available data. We have selected a consistent 25-micron squared field of view.

Here we detail how we test the robustness of our model against synthetic data. For the field of view, we choose 25-micron squared as this best represents our experimental data. We then proceed to simulate particle trajectories, with varying diffusion maps, using Equation (1).

We begin with a flat plane, as a preliminary sanity check, and then progressively add more variation to the ground truth surface to see where the inference breaks down. We also check how the relative error with the ground truth changes with varying amounts of data, ranging from 5 *×* 10^3^ to 2 *×* 10^5^ positions. We continue to use 10% relative error, Equation (13), as the benchmark for accurate inference. As can be seen, resolving a higher variation in diffusion coefficient requires more information (data), but at 2 *×* 10^5^ positions, we resolve a diffusion coefficient map consisting of waves with a maximum period of 40% of the field of view.

### S0.2 Consideration of Temporal Changes in Diffusion

To illustrate that there are minimal temporal changes in the diffusion maps of our experimental data or that different trajectories show minimal diffusion coefficient variation when passing through the same region of space, we consider a self-consistency check where we split the data in half by randomly selecting trajectories. As trajectories pass through different points in space at different times, obtaining consistent diffusion maps from different trajectories would illustrate that the diffusion maps are stable in time.

We test this by considering a dataset that is very dense with trajectories. Of the experimental datasets considered in the paper, Dataset 1 is populated with the most trajectories and localizations, thus this is the dataset we consider. Furthermore, to minimize errors due to areas of no data we zoom into a 10-micron square that has very few pockets of no data. Figure S2 shows the results of this analysis. In particular, we find that, in regions close to data we have less than 10% relative error.

**Figure S2:**
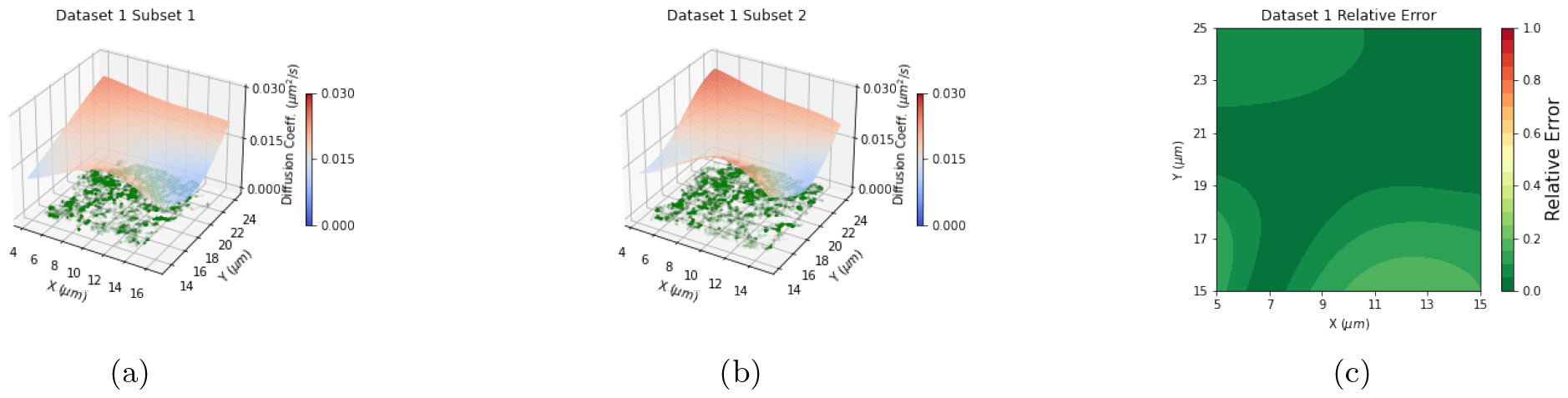
Self-consistency by subsetting trajectories. Here we zoom to the most trajectory-rich region of our first experimental dataset. We subset the data in this region by randomly splitting the trajectories into two groups. Then we run our inference scheme on each subset individually, (a) and (b). Then we compare by plotting relative error, (c).

### S0.3 Comparison with Previous Results

As shown in previous work [29], which perform identical experiments on similar CHO cells, we expect a diffusion coefficient magnitude broadly centered at about 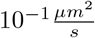to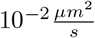. To compare, for all six experimental datasets, we histogram the estimated diffusion at all positions with protein localization, Figure S3. Our results are consistent, with most of the density representing a diffusion coefficient neighboring the expected 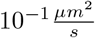to 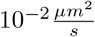 magnitude.

**Figure S3:**
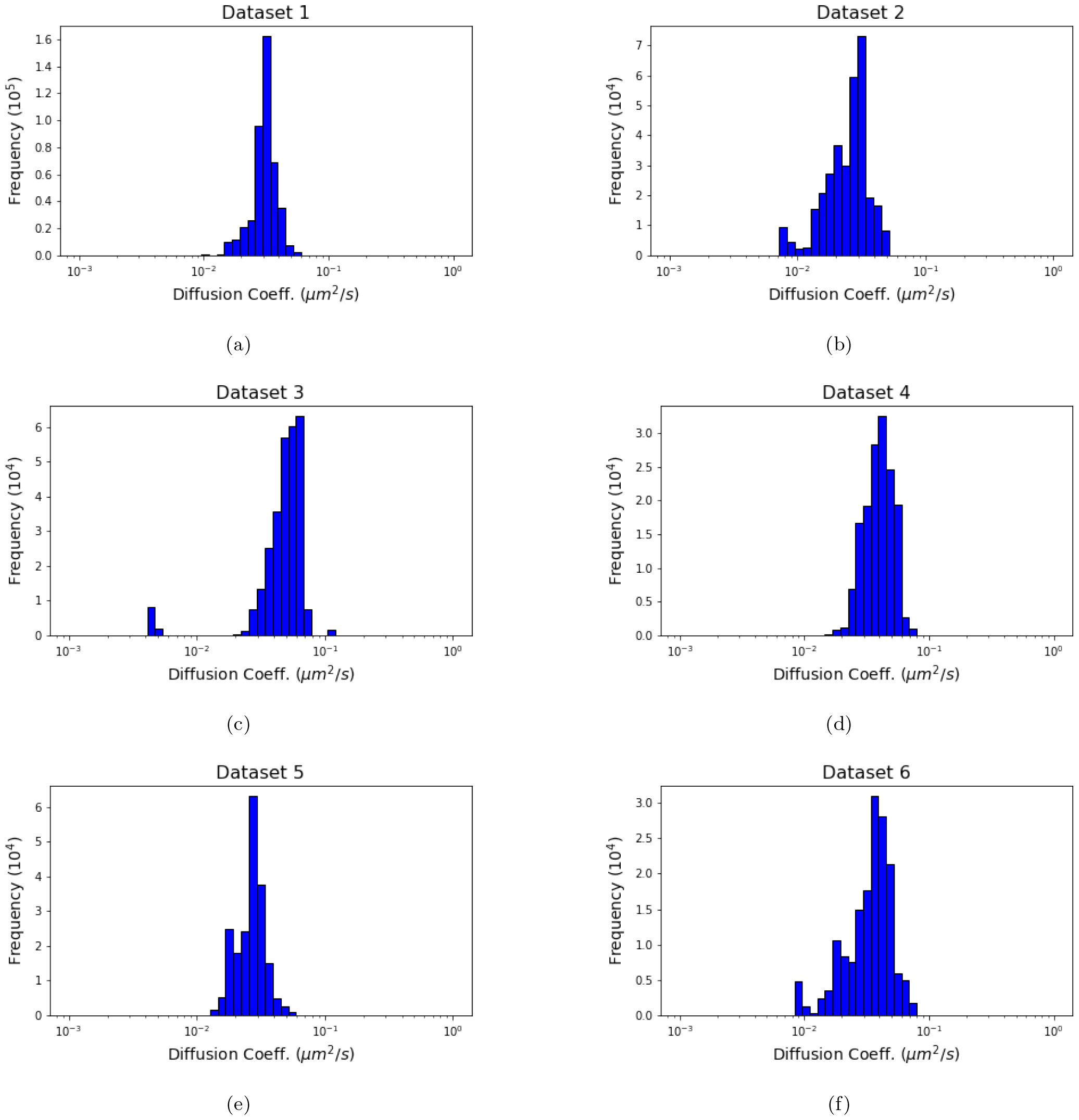
Histogram of estimated diffusion rates. For each of the six experimental datasets considered throughout, we visualize the frequency of estimated diffusion coefficients at all positions of every trajectory. Specifically, we use the inferred diffusion maps shown in Figure 3 to determine the diffusion values at all protein localizations in the data. This evaluation is done using the interpolation scheme defined in Equation (9).

## Notes

### Competing Interest Statement

SP, SB, and VK are affiliated with Saguaro AI LLC.

## References

[1] Xiaolin Cheng and Jeremy C. Smith. Biological Membrane Organization and Cellular Signaling. Chemical Reviews, 119(9):5849–5880, 2019.

[2] Doralicia Casares, Pablo V. Escribá, and Catalina Ana RossellÓ. Membrane Lipid Composition: Effect on Membrane and Organelle Structure, Function and Compartmentalization and Therapeutic Avenues. International Journal of Molecular Sciences, 20(9):2167, 2019.

[3] György Vereb, János Szöllosi, János Matkó, Peter Nagy, Tamás Farkas, László Vigh, László Mátyus, Thomas A. Waldmann, and Sándor Damjanovich. Dynamic, yet structured: The cell membrane three decades after the Singer-Nicolson model. Proceedings of the National Academy of Sciences of the United States of America, 100(14):8053–8058, 2003.

[4] Ken Jacobson, Ping Liu, and B Christoffer Lagerholm. The Lateral Organization and Mobility of Plasma Membrane Components. Cell, 177(4):806–819, 2019.

[5] Tatsunari Ohkubo, Takaaki Shiina, Kayoko Kawaguchi, Daisuke Sasaki, Rena Inamasu, Yue Yang, Zhuoqi Li, Keizaburo Taninaka, Masaki Sakaguchi, Shoko Fujimura, Hiroshi Sekiguchi, Masahiro Kuramochi, Tatsuya Arai, Sakae Tsuda, Yuji C. Sasaki, and Kazuhiro Mio. Visualizing Intramolecular Dynamics of Membrane Proteins. International Journal of Molecular Sciences, 23(23):14539, 2022.

[6] Rumana Mahtarin, Shafiqul Islam, Md. Jahirul Islam, M. Obayed Ullah, Md. Ackas Ali, and Mohammad A. Halim. Structure and dynamics of membrane protein in SARS-CoV-2. Journal of Biomolecular Structure and Dynamics, 40(10):4725–4738, 2022.

[7] Konstantinos Tsekouras, Amanda P. Siegel, Richard N. Day, and Steve Pressé. Inferring Diffusion Dynamics from FCS in Heterogeneous Nuclear Environments. Biophysical Journal, 109(1):7–17, 2015.

[8] Jean-Baptiste Masson, Patrice Dionne, Charlotte Salvatico, Marianne Renner, Christian G Specht, Antoine Triller, and Maxime Dahan. Mapping the energy and diffusion landscapes of membrane proteins at the cell surface using high-density single-molecule imaging and Bayesian inference: application to the multiscale dynamics of glycine receptors in the neuronal membrane. Biophysical journal, 106(1):74–83, 2014.

[9] Suman Ranjit, Enrico Gratton, and Luca Lanzano. Mapping Diffusion in a Living Cell using the Phasor Approach. Biophysical Journal, 107(12):2775–2785, 2014.

[10] Dominique Ernst and Jürgen Köhler. Measuring a diffusion coefficient by single-particle tracking: statistical analysis of experimental mean squared displacement curves. Physical Chemistry Chemical Physics, 15(3):845– 849, 2013.

[11] Laura Toppozini, Victoria Garcia-Sakai, Robert Bewley, Robert Dalgliesh, Toby Perring, and Maikel C Rheinstädter. Diffusion in membranes: Toward a two-dimensional diffusion map. EPJ Web of Conferences, 83:02019, 2015.

[12] Yerim Lee, Carey Phelps, Tao Huang, Barmak Mostofian, Lei Wu, Ying Zhang, Kai Tao, Young Hwan Chang, Philip JS Stork, Joe W Gray, et al. High-throughput, single-particle tracking reveals nested membrane domains that dictate KRasG12D diffusion and trafficking. eLfe, 8:e46393, 2019.

[13] Silvan Türkcan, Antigoni Alexandrou, and Jean-Baptiste Masson. A Bayesian Inference Scheme to Extract Diffusivity and Potential Fields from Confined Single-Molecule Trajectories. Biophysical Journal, 102(10):2288– 2298, 2012.

[14] Jesús Pineda, Benjamin Midtvedt, Harshith Bachimanchi, Sergio Noé, Daniel Midtvedt, Giovanni Volpe, and Carlo Manzo. Geometric deep learning reveals the spatiotemporal features of microscopic motion. Nature Machine Intelligence, 5(1):71–82, 2023.

[15] Mohamed El Beheiry, Maxime Dahan, and Jean-Baptiste Masson. InferenceMAP: mapping of single-molecule dynamics with Bayesian inference. Nature Methods, 12(7):594–595, 2015.

[16] Mary A Rohrdanz, Wenwei Zheng, Mauro Maggioni, and Cecilia Clementi. Determination of reaction coordinates via locally scaled diffusion map. The Journal of Chemical Physics, 134(12):03B624, 2011.

[17] Andrew Gordon Wilson and Hannes Nickisch. Kernel Interpolation for Scalable Structured Gaussian Processes (KISS-GP). CoRR, abs/1503.01057, 2015.

[18] Steve Pressé and Ioannis Sgouralis. Data Modeling for the Sciences: Applications, Basics, Computations. Cambridge University Press, 2023.

[19] Carl Edward Rasmussen and Christopher K. I. Williams. Gaussian Processes for Machine Learning. The MIT Press, 2005.

[20] Robert Zwanzig. Nonequilibrium statistical mechanics. Oxford University Press, 2001.

[21] Issei Sato and Hiroshi Nakagawa. Approximation analysis of stochastic gradient Langevin dynamics by using Fokker-Planck equation and Ito process. In International Conference on Machine Learning, pages 982–990. PMLR, 2014.

[22] M. E. Young, P. A. Carroad, and R. L. Bell. Estimation of diffusion coefficients of proteins. Biotechnology and Bioengineering, 22(5):947–955, 1980.

[23] Michalis K Titsias, Neil Lawrence, and Magnus Rattray. Markov chain Monte Carlo algorithms for Gaussian processes. Inference and Estimation in Probabilistic Time-Series Models, 9:298, 2008.

[24] Malcolm Sambridge. A Parallel Tempering algorithm for probabilistic sampling and multimodal optimization. Geophysical Journal International, 196(1):357–374, 2013.

[25] Shalini T Low-Nam, Keith A Lidke, Patrick J Cutler, Rob C Roovers, Paul MP van Bergen en Henegouwen, Bridget S Wilson, and Diane S Lidke. ErbB1 dimerization is promoted by domain co-confinement and stabilized by ligand binding. Nature Structural & Molecular Biology, 18(11):1244–1249, 2011.

[26] Khuloud Jaqaman, Dinah Loerke, Marcel Mettlen, Hirotaka Kuwata, Sergio Grinstein, Sandra L Schmid, and Gaudenz Danuser. Robust single-particle tracking in live-cell time-lapse sequences. Nature Methods, 5(8):695– 702, 2008.

[27] Melissa Linkert, Curtis T Rueden, Chris Allan, Jean-Marie Burel, Will Moore, Andrew Patterson, Brian Loranger, Josh Moore, Carlos Neves, Donald MacDonald, et al. Metadata matters: access to image data in the real world. Journal of Cell Biology, 189(5):777–782, 2010.

[28] Juan A Torreno-Pina, Bruno M Castro, Carlo Manzo, Sonja I Buschow, Alessandra Cambi, and Maria F Garcia-Parajo. Enhanced receptor–clathrin interactions induced by N-glycan–mediated membrane micropatterning. Proceedings of the National Academy of Sciences, 111(30):11037–11042, 2014.

[29] Carlo Manzo, Juan A Torreno-Pina, Pietro Massignan, Gerald J Lapeyre Jr, Maciej Lewenstein, and Maria F Garcia Parajo. Weak ergodicity breaking of receptor motion in living cells stemming from random diffusivity. Physical Review X, 5(1):011021, 2015.

[30] IV Bryan, J. Shepard, Ioannis Sgouralis, and Steve Pressé. Inferring effective forces for Langevin dynamics using Gaussian processes. The Journal of Chemical Physics, 152(12):124106, 2020.

[31] Ioannis Sgouralis, Lance W.Q. Xu, Ameya P. Jalihal, Nils G. Walter, and Steve Pressé. BNP-Track: A framework for superresolved tracking. bioRxiv, 2023.

[32] Sina Jazani, Ioannis Sgouralis, and Steve Pressé. A method for single molecule tracking using a conventional single-focus confocal setup. The Journal of Chemical Physics, 150(11):114108, 2019.

[33] Katharina Gaus, Enrico Gratton, Eleanor P. W. Kable, Allan S. Jones, Ingrid Gelissen, Leonard Kritharides, and Wendy Jessup. Visualizing lipid structure and raft domains in living cells with two-photon microscopy. Proceedings of the National Academy of Sciences, 100(26):15554–15559, 2003.

[34] Sjövall, Peter and Lausmaa, Jukka and Nygren, Håkan and Carlsson, Lennart and Malmberg, Per. Imaging of membrane lipids in single cells by imprint-imaging time-of-flight secondary ion mass spectrometry. Analytical Chemistry, 75(14):3429–3434, 2003.

[35] Ingólfsson, Helgi I. and Melo, Manuel N. and van Eerden, Floris J. and Arnarez, Clément and Lopez, Cesar A. and Wassenaar, Tsjerk A. and Periole, Xavier and de Vries, Alex H. and Tieleman, D. Peter and Marrink, Siewert J. Lipid organization of the plasma membrane. Journal of the American Chemical Society, 136(41):14554–14559, 2014.

